# Low protein diet protects liver function upon Salmonella infection by metabolic reprogramming of macrophages

**DOI:** 10.1101/2024.03.01.582753

**Authors:** Edyta E Wojtowicz, Katherine Hampton, Mar Moreno-Gonzalez, Charlotte L Utting, Yuxuan Lan, Paula Ruiz, Gemma Beasy, Caitlin Bone, Charlotte Hellmich, Rebecca Maynard, Luke Acton, Andrea Telatin, Robert A Kingsley, Iain C Macaulay, Stuart A Rushworth, Naiara Beraza

**Author notes:** **Corresponding Authors:** To whom correspondence should be addressed. Naiara Beraza, PhD Gut Microbes and Health institute strategic programme, Food Microbiome and Health institute strategic programme, Quadram Institute, Norwich Research Park, Norwich, UK., Stuart A Rushworth, PhD Centre for Metabolic Health, Faculty of Medicine, University of East Anglia, Norwich Research Park, Norwich, UK.

## Abstract

**Background & Aims:** Western diets are the underlying cause of metabolic and liver diseases. Recent trend to limit the consumption of protein-rich animal products has become more prominent. This dietary change entails decreased protein consumption; however, it is still unknown how this affects innate immunity. Here, we studied the influence of a low protein diet (LPD) on the liver response to bacterial infection.

**Methods:** Mice were fed a LPD and exposed to *Salmonella enterica* serotype Typhimurium infection. Mechanistic studies were done *in vitro* where bone marrow derived macrophages were cultured in a low-aa media to mimic *in vivo* reduction of protein availability and challenged with bacterial endotoxin.

**Results:** We found that a LPD protects from *S* Typhimurium-induced liver damage. Bulk- and 10xsingle cell-RNA sequencing of liver tissues and isolated immune cells showed reduced activation of myeloid cells in mice fed with LPD after *S* Typhimurium infection. Mechanistically, we found reduced activation of the mammalian target of rapamycin (mTOR) pathway whilst increased phagocytosis and activation of autophagy in LPD-programmed macrophages. Dietary restoration of leucine reverted the protective effects of a LPD and restored the damaging effects of Salmonella on liver parenchyma in mice.

**Conclusions:** Low protein diet protects the liver form *S* Typhimurium-induced tissue damage via modulating macrophage autophagy and phagocytosis. Our result support the causal role of dietary components on the fitness of the immune system.

**SYNOPSIS:** Low protein diet protects the liver from Salmonella-mediated liver injury that associates with reduced mTOR activation and increased autophagy in macrophages. Restoration of the mTOR pathway with aminoacid supplementation reverses the protection of a low protein diet from Salmonella-liver damage.

---

Metabolism and immunity share a close connection at both the cellular and organismal levels. Metabolic regulation of the innate and adaptive immune responses is an active and expanding area of investigation. In particular, the roles of dietary choices in regulating the functions of different immune cells[1] and as the underlying cause of chronic diseases[2] are increasingly recognized.

Western diets rich in processed foods are linked to metabolic diseases. They also cause altered composition and functional states of various immune cells in tissues, contributing to chronic activation of macrophages and inflammation[3–5]. Epidemiological studies have revealed an increased susceptibility to infections in patients with diabetes or obesity, pointing to an evident dysfunction in the immune response[6, 7].

On the opposite side of the Western-diet spectrum, a part of the population is adopting healthier dietary habits that include reduced consumption of meat products and increased consumption of plant-based products. This nutritional adjustment encompasses significantly reduced intake of protein[8]. Protein malnutrition (0.5-2.5% protein) has adverse consequences in young children and juvenile mice for the immune system function[9–12], leading to immunosuppression[11, 12], while a reduction in dietary protein (7-10% protein) inhibits cancer development[13, 14] and metastasis[15]. Still, our knowledge on whether the physiological reduction of protein intake impacts host immunity and response to infection is limited.

To rapidly identify pathogens, cells of the innate immune system including macrophages which engulf bacteria and traffic the resulting phagosomes through the fusion with lysosomes that results in their destruction[16]. Additionally, macrophages secrete proinflammatory and antimicrobial mediators to inactivate pathogens[17, 18]. Bacteria, including *Salmonella enterica* serotype Typhimurium (*S*. Typhimurium), have developed mechanisms to escape innate immunity through triggering macrophages necroptosis[19, 20] or circumventing autophagy by directing for degradation molecular sensors-AMPK and SIRT1[21]. Both AMPK and SIRT1 regulate the evolutionary conserved nutrient sensing pathway-mammalian target of rapamycin (mTOR)[22]. mTOR integrates the metabolic, autophagic and phagocytic state of the cell, therefore linking the innate immunity with the host metabolic state[23].

Dietary nutrients, more specifically amino acids (in particular serine, glutamine, leucine and arginine) regulate innate immunity, specifically they shift the balance between proinflammatory (M1-like) or pro-healing (M2-like)[24] macrophage subsets. Glutamine and serine depletion promotes proinflammatory M1-like macrophages producing IL-1b[25–27], while the inhibition of serine synthesis decreases IL-1b and TNF production in LPS-induced endotoxemia[28]. Arginine depletion induces M2-like macrophages facilitating proliferation and healing[29]. Leucine abundance activates the metabolic master regulator mTOR (specifically the mTORC1 subunit) by providing the acetyl group[30], enhancing glycolysis and proinflammatory M1 macrophage state. It leads to the inflammasome activation, release of proinflammatory cytokines[31], while shutting down autophagy, thus promoting survival of pathogens like *S*. Typhimurium within macrophages[32]. The expression of the leucine transporter-Slc7a5 also underlies macrophage metabolic rewiring by limiting the cellular levels of leucine. Lipopolysaccharide (LPS) increases leucine-transporter expression in macrophages, while its pharmacological inhibition reduces glycolysis and IL-1b production[33].

Despite this body of evidence on the key role of amino acids for fine tuning immune response it is unknown how a systemic lower abundance of dietary amino acids regulates host innate immune response to pathogen infection.

In the present study, we test the hypothesis that a LPD effects the host liver response to *S*. Typhimurium infection by the transcriptional and functional reprogramming of macrophages in the liver. Here, we provide a mechanistic insight into the beneficial role of a LPD on bacteria clearance and preserving liver function after infection.

## RESULTS

### Low protein diet protects liver function from *S*. Typhimurium induced damage

To evaluate the impact of reduced protein intake on the liver response to infection we fed C57/Bl6 mice *ad libitum,* a normal (control) or a low protein (LPD) isocaloric diets for 10 weeks after which mice were inoculated with *S*. Typhimurium (Fig 1A). Three days post inoculation we observed a significant reduction of circulating alanine transaminase (ALT) in plasma of LPD-fed mice after *S*. Typhimurium infection (8-fold decrease) suggesting decreased liver injury compared to normal diet fed animals (Fig. 1B). Histopathological analysis of liver sections with hematoxylin and eosin (H&E) staining revealed reduced areas of necrosis consistent with reduced liver injury in LPD fed mice compared to normal diet after *S*. Typhimurium infection (Fig. 1C).

**Figure 1.**
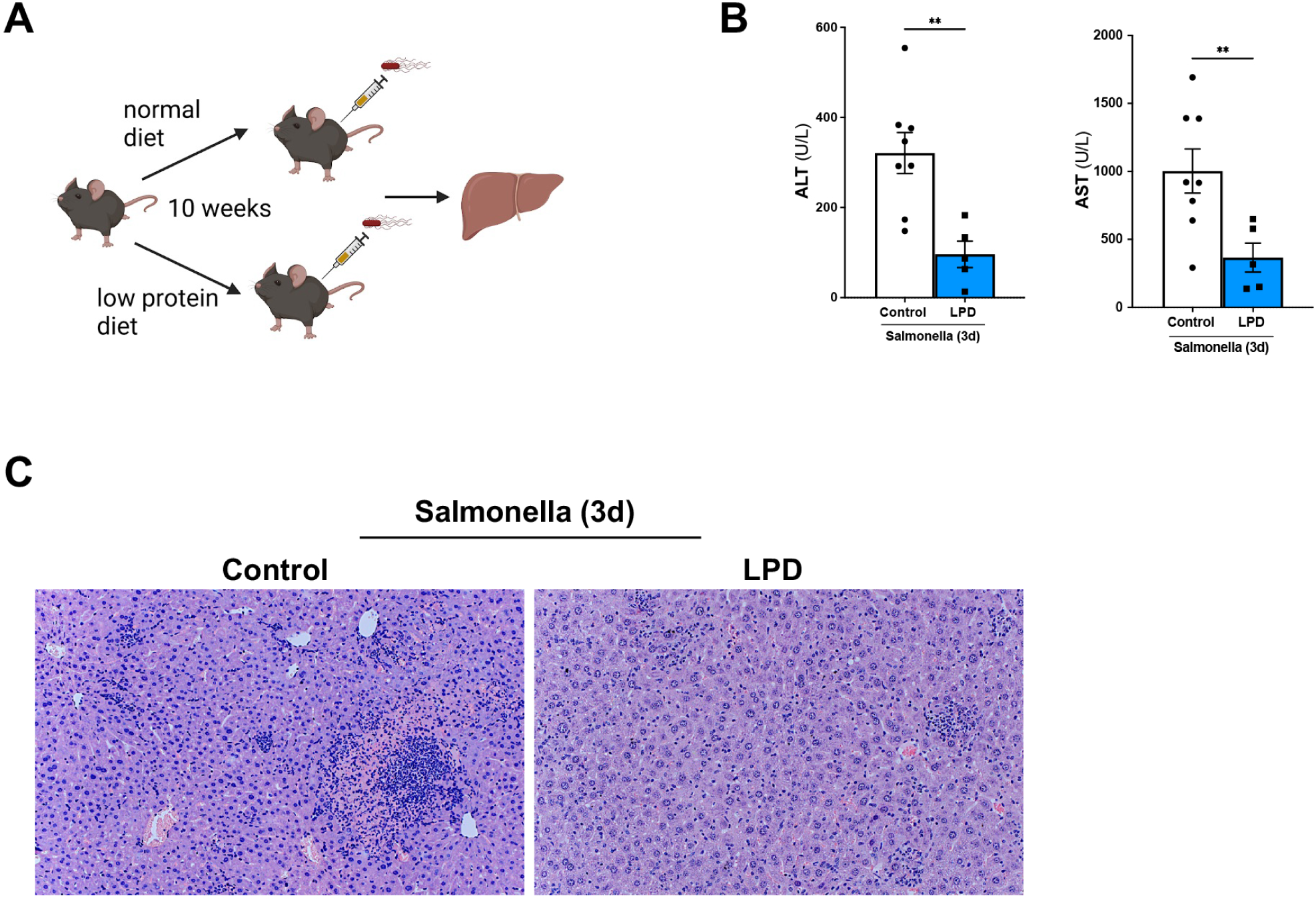
LPD limits *S.* Typhimurium induced liver damage in vivo. **(A)** Experimental set up scheme**. (B)** Quantification of the serum levels of alanine aminotransferase (ALT) and aspartate aminotransferase (AST) in animals fed with a control or low protein diet (LPD) for 10 weeks and 3 days after *S.* Typhimurium infection. **(C)** H&E staining in liver sections obtained from control and LPD fed mice infected with *S.* Typhimurium. Arrows point to necrotic areas. Analyses were done from n=5-8 mice. Results shown are representative from 3 independent experiments. Representative microscopic images are shown from 20x magnification. Values are mean ± SEM. \*\**P* <0.01 (Control diet vs LPD).

### LPD attenuates expression of proinflammatory cytokines and chemokines in liver cells after *S.* Typhimurium infection

To better understand the molecular changes in livers from normal diet and LPD fed animals after *S*. Typhimurium infection we performed bulk RNA-seq was conducted on liver lysates from mice given normal or LPD diet for 10 weeks and subsequently infected with *S.* Typhimurium for 3 days. The liver transcriptional profile of infected mice on the LPD clustered distinctly compared to that of infected mice on a normal diet by PCA (Fig. 2A) and the heatmap (Fig. 2B). The analysis of differentially expressed genes (FDR, q<0.001) between normal and LPD diets after Salmonella infection showed: 2470 up-regulated genes and 2026 down-regulated genes in mice fed with a normal diet, while LPD feeding led to 2301 up-regulated and 1860 down-regulated genes after Salmonella.

**Figure 2.**
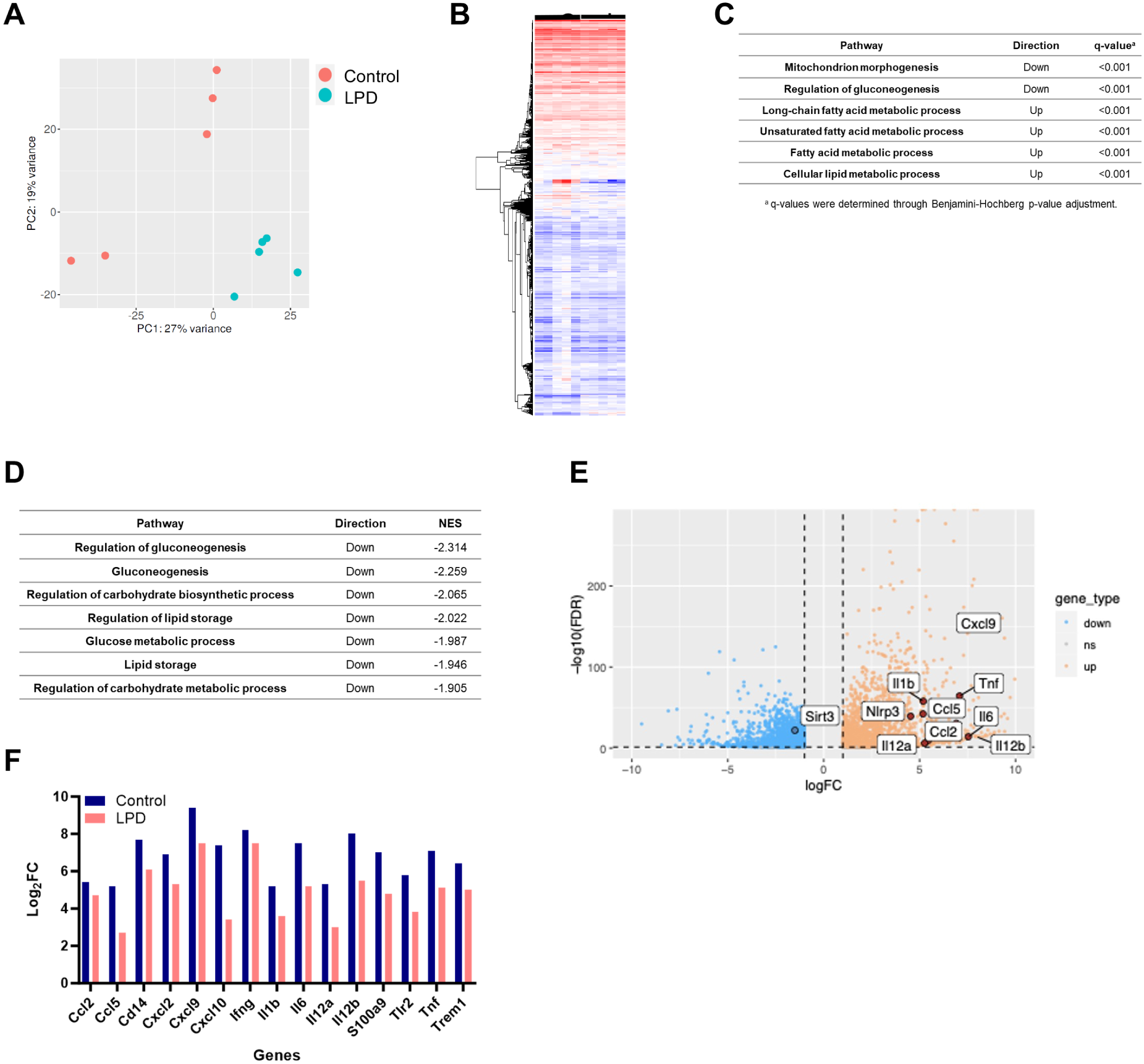
Bulk RNA sequencing of liver samples showing diet induced metabolic reprogramming and anti-inflammatory changes. **(A)** Principal component analysis (PCA) plot on raw gene count data for first and second component in control and LPD liver samples infected with *S.* Typhimurium. **(B)** Heatmap of raw gene count data between treatment groups. **(C)** Table including the enrichment analysis using Gene Ontology (GO) database. **(D)** Table including pathway analysis using GO, X-axis represents the normalized enrichment score (NES), y-axis the gene sets. An increase in the gene set shows positive NES, and negative NES represents a decrease in the gene set. **(E)** Volcano plot of gene expression from control and LPD fed animals. The blue dots represent down-regulated differentially expressed genes; orange dots represent the upregulated differentially expressed genes. **(F)** Fold change (Log_2_FC) of immune-related genes in control and LPD. Immune related genes are down-regulated in LPD diet compared to control diet. All q-value were <0.001; q-values were determined through Benjamini-Hochberg p-value adjustment.

The enrichment analysis indicated that LPD induced molecular changes in liver cells compared to normal diet in mice after infection (Suppl. Fig. 1 Table1). From the gene set enrichment analysis (GSEA), down-regulated key metabolic pathways included gluconeogenesis, lipid metabolism and OXPHOS-related processes in LPD samples (Fig. 2C). proinflammatory related genes were down-regulated in LPD compared to normal diet after infection including *Cxcl2, Tnf, Tlr2, Nlrp3, Il1b, Il6, Il12a* (Figure 2D-F) pointing to immunoregulatory effects of LPD feeding. In addition, we observed changes in the expression pattern for *Sirt3* which was upregulated in LPD compared to normal diet cells after *S.* Typhimurium infection, suggesting further metabolic changes including the modulation of autophagy[34] and inflammation[35].

Together, our results point to metabolic rewiring and anti-inflammatory effects of LPD on the liver upon *S.* Typhimurium infection.

### Single cells analysis of immune cells in the liver captures reprogramming of monocytes

To identify changes in cellular composition and gene expression programs specifically in innate immune cells, we performed single cell RNA sequencing using the 10X Chromium platform. We isolated immune cells from livers[36], followed by FACS purification for viable, CD45^+^ cells from control (n=3) and LPD (n=3) mice at 3 days post *S.* Typhimurium infection. We generated standard 10X 3’ scRNA-seq libraries from FACS-purified cell populations from control and LPD samples after infection (Fig. 3A).

**Figure 3.**
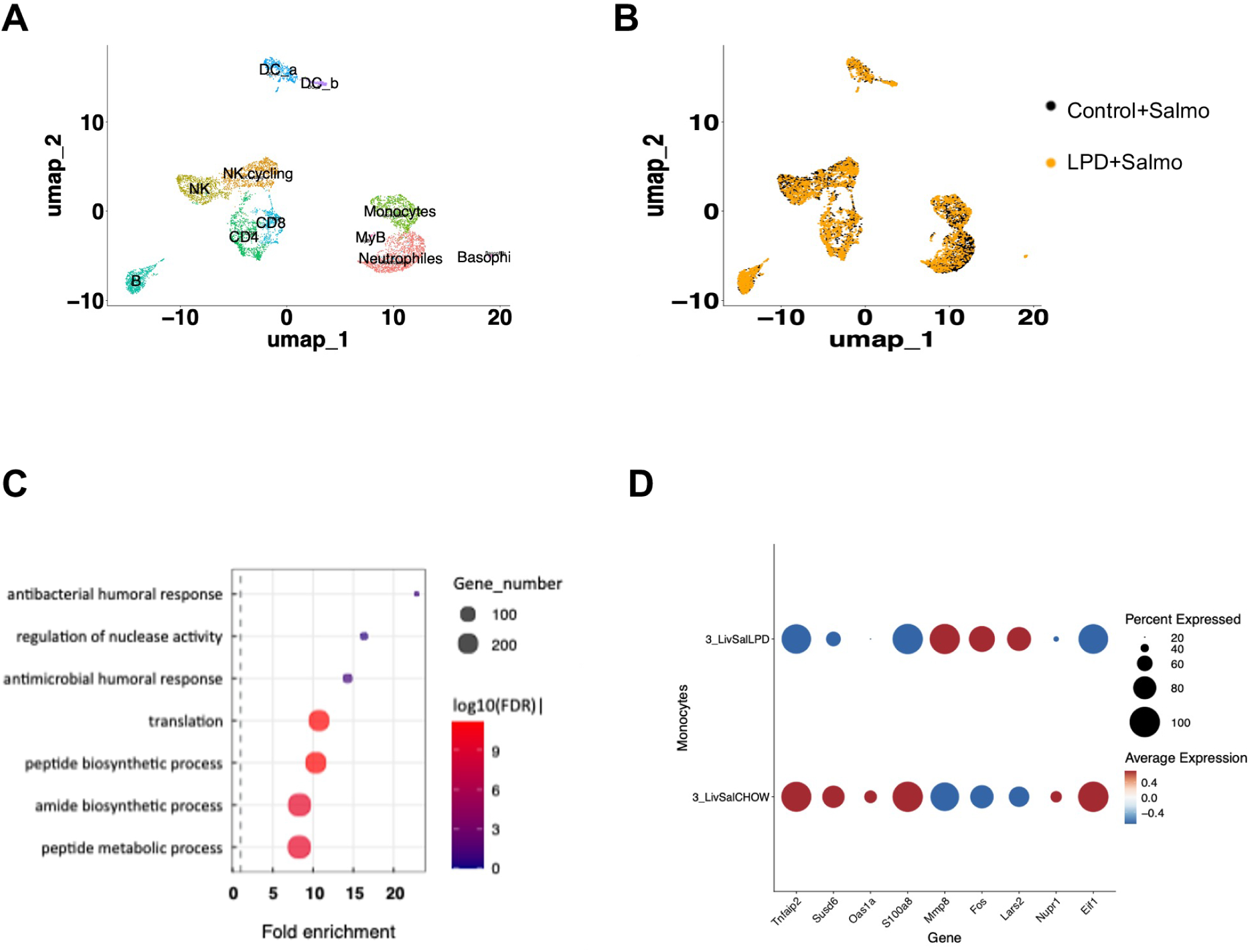
Single cell RNA sequencing on immune cells isolated from livers after infection. **(A)** UMAP representing cell types present among liver immune cells in control and LPD fed animals after *S.* Typhimurium infection. **(B)** UMAP representing the cell type distribution in samples from animals fed control and LPD diets. **(C)** KEGG analysis of pathways enriched in control and LPD fed diets highlighting the metabolic remodelling of monocytes and enrichment for innate immune response genes in LPD derived monocytes. **(D)** Dot plot representing differentially expressed genes between monocytes from control and LPD diet.

This analysis identified 11 clusters within CD45^+^ liver cell population, which can be manually annotated using RNA marker expression (Suppl. Figure 1). We did not observe a significant shift in the frequency of analysed cell types (Fig. 3B).

Differential gene expression analysis in control and LPD liver monocytes after *S.* Typhimurium revealed enrichment for nuclease activity, humoral and antibacterial response in normal diet compared to LPD monocytes after infection (Fig. 3C). We detected 53 downregulated and 21 upregulated genes in monocytes (Fig. 3D). Thus, after infection, monocytes from LPD-fed mice had decreased levels of *Tnfaip2* (Fig. 3D), which expression is regulated by TNFα and other proinflammatory stimuli like IL1β and LPS via NFκB activation[37]. Our results also showed a reduction of *Susd6* (Fig. 3D) that promotes chemokine expression[38]. Furthermore, liver monocytes from LPD-fed mice expressed decreased levels of *Oas1a* (Fig. 3D), which stimulates the expression of chemokines (Ccl2, Ccl3, Ccl4, Ccl8, Cxcl9 and Cxcl10) upon inflammation in macrophages[39]. Additionally, LPD liver-monocytes expressed decreased levels of *S100a8* (Fig. 3D), which increases recruitment of leukocytes producing proinflammatory cytokines[40]. Importantly, liver monocytes from LPD-fed mice expressed increased levels of *Mmp8* (Fig. 3D), a known inhibitor of macrophage *Mip-1α*, which drives the acute lung inflammation in mice[41]. Monocytes from infected mice fed with LPD expressed significantly higher *c-Fos* levels compared to control diet-fed animals (Fig. 3D), a transcription factor involved in modulation of inflammation and susceptibility to Salmonella infection[42, 43].

Interestingly, we observed decreased levels of stress-induced transcription factor *Nupr1* (Fig. 3D), which activates mTOR pathway[44]. In line with this, we also found decreased expression of EIf1 that forms the 48S complex with mTOR effector Eif2 mediating its activation[45, 46], in liver monocytes from LPD-fed mice after Salmonella.

Together, our results point to a less proinflammatory phenotype of these cells along with modulation of metabolic factors, including mTOR signaling.

### Low amino acid availability decreases the expression of inflammasome components, proinflammatory cytokines and mTOR activation in bone marrow derived macrophages

Nutrient sensing pathway-mTOR is the key regulator of monocyte/macrophage response to infection by promoting inflammasome activation and thus production of IL1β proinflammatory cytokine responses while restricting the activation of autophagy[47]. mTOR is regulated by amino acid availability[48] and this balance may mediate the effect of the LPD on the macrophage activation we observed in response to Salmonella infection *in vivo*.

To test the hypothesis that changes in amino acid availability affects macrophage activation, we performed *in vitro* experiments using bone marrow derived macrophages (BMDMs) that we cultured in a low amino acid media (herein low-aa) to mimic LPD condition *in vivo*. As a model of infection, we stimulated BMDM cultured in control or low-aa media with lipopolysaccharide (LPS) for the indicated times (Fig. 4A). To measure the activation of the mTOR pathway we performed intracellular FACS staining for phosphorylated S6 kinase (pS6K, Ser 235, 236) and observed 1.5-fold increased levels of pS6K in control compared to low-aa BMDMs (Fig. 4B, C), confirming reduced mTOR activation in the latter.

**Figure 4.**
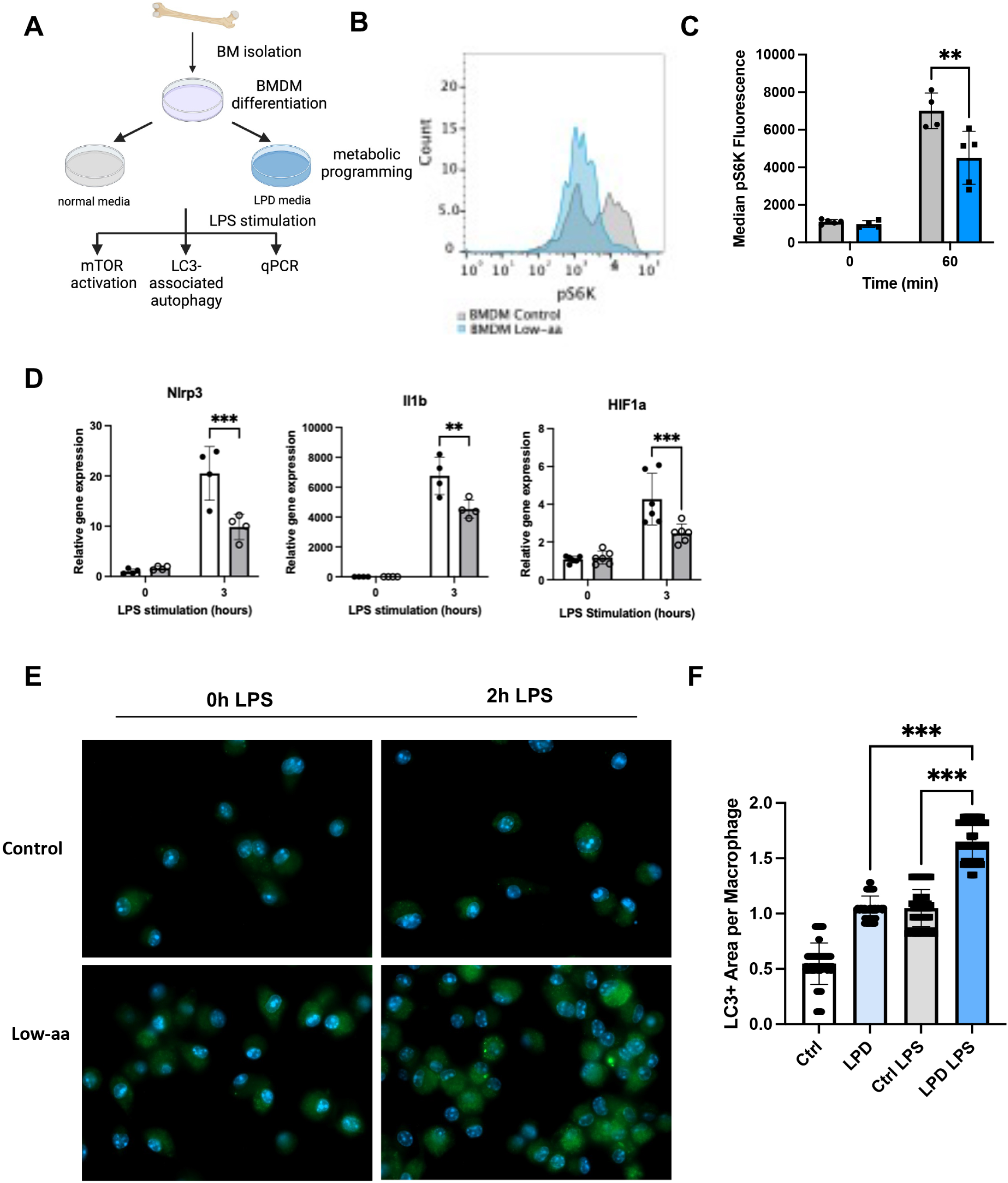
Metabolic reprogramming of In vitro derived BMDM’s inhibits mTOR and activates LC-3 dependent autophagy. **(A)** Experimental set up. **(B)** Representative histogram depicting decreased MFI of pS6 kinase level in Low-aa media programmed BMDM’s compared to normal media. **(C)** Quantification of pS6 kinase level upon LPS treatment of BMDM’s from normal or Low-aa media. **(D)** qPCR expression analysis of Nlrp3, Il1b and HIF1a in metabolically reprogrammed BMDM. (**E**) Representative images of immunofluorescence staining for LC3 to quantify the degree of autophagy in reprogrammed BMDM’s. (**F**) Quantification of the autophagy level in normal and LPD programmed BMDM’s, Representative images are shown from 63x magnification. *In vitro* experiments were repeated 2-3x with n=3-4 replicates. Values are mean ± SEM. \*\**P* <0.01, \*\*\**P* <0.001.

Next, we determined the levels of the inflammasome subunit Nlrp3, which was 2-times lower expressed in low-aa BMDMs compared to control after LPS stimulation. Reduced Nlrp3 levels correlated with decreased expression of IL1β (Fig. 4D) and decreased Hif1α expression (Fig. 4D) after LPS, supporting the reduced activation of macrophages in a low-aa media.

The cross-regulation between inflammasome and autophagy to control the inflammatory response is well established[47]. Thus, we next evaluated the level of autophagy in BMDMs exposed to low-aa media by detecting LC3, a surrogate of autophagy. Immunofluorescence staining of LC3 puncta and its further quantification revealed increased autophagy activity in BMDMs cultured in low-aa compared to control media in response to LPS (Fig. 4E, F).

Together these results show that low-aa availability, consistent with a low protein diet feeding *in vivo*, reprograms BMDMs to become less proinflammatory while increasing autophagy.

### Restoration of mTOR activation with dietary leucine supplementation abolishes the LPD-mediated protective effects on liver function via regulating macrophage phagocytosis and autophagy

Next, we aimed to assessing the physiological impact of metabolic reprogramming in macrophages via the re-activation of mTOR in response to infection *in vivo*. To do this, we supplemented the LPD with Leucine, an essential amino acid well-characterised as an activator of the mTOR pathway[49]. A group of mice was fed with LPD diet for 7 weeks and then switched to LPD diet supplemented with 3% leucine (LPD+Leu) while another group was fed with a LPD for the duration of the experiment (10w) as a control (Fig. 5A).

**Figure 5.**
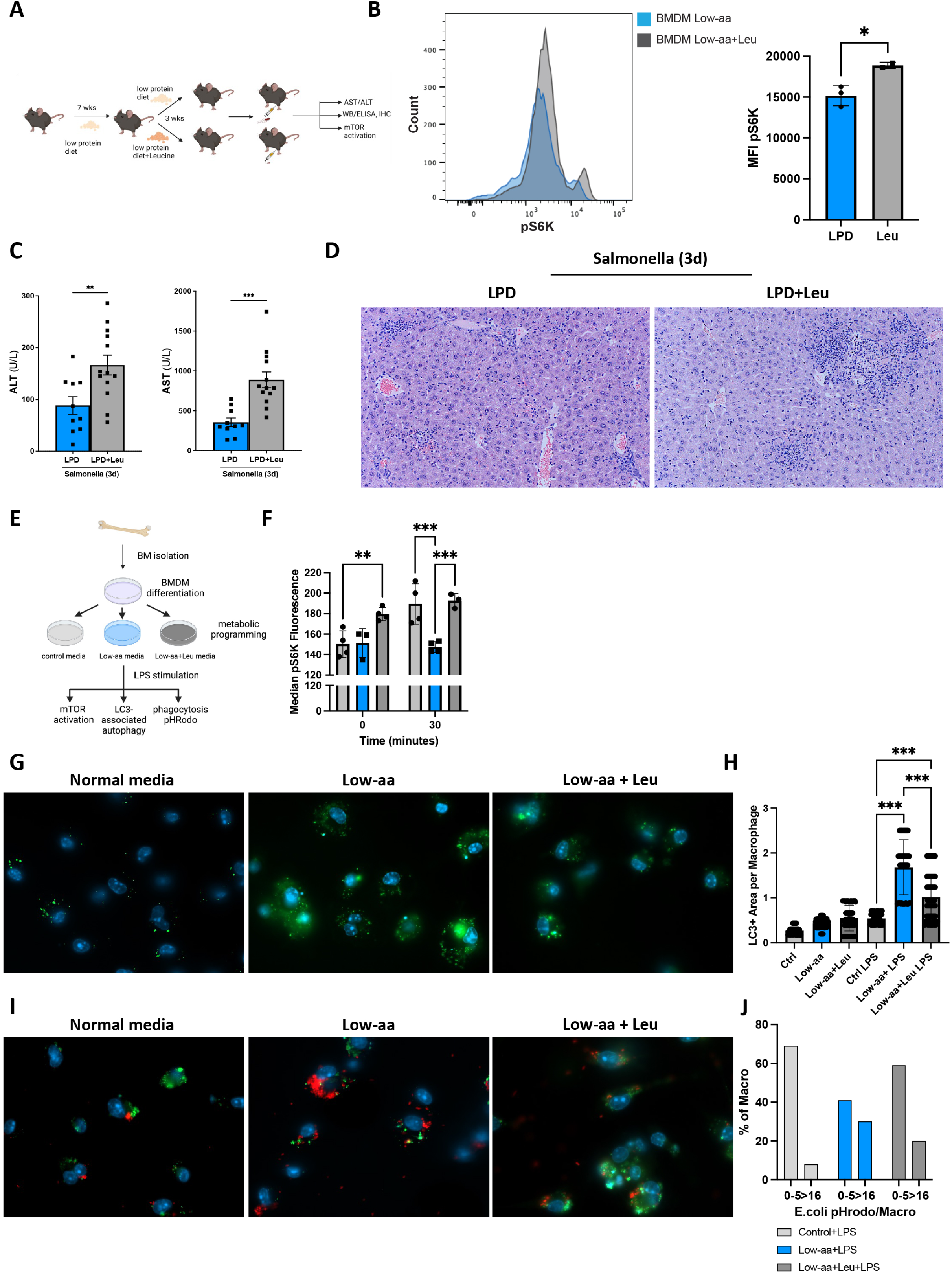
Supplementation with leucine diminishes LPD-mediated protection from liver damage upon *S*. Typhimurium infection via BMDM metabolic reprogramming. **(A)** In vivo experimental set up. **(B)** Intracellular FACS staining for pS6 kinase in primary bone marrow F4/80+ macrophages stimulated 60 min with LPS and quantification of pS6 kinase MFI. **(C) Serum** level of liver enzymes ALT and AST in animals infected with S.Typhimurium. **(D)** H&E staining on liver sections from LPD and LPD+Leu fed mice, 3 days after S.Typhimurium infection. **(E)** In vitro experimental set up for BMDM. **(F)** Quantification of intracellular FACS staining for pS6 kinase in metabolically programmed BMDM’s with normal, Low-aa or Low-aa+Leu media. **(G)** Immunofluorescence staining for LC3 evaluating autophagy in metabolically reprogrammed BMDM’s and **(H)** further quantification of immunofluorescence. **(I)** Immunofluorescence for autophagy associated LC3-in green and pHrodo E. coli beads in lysosome depicted in red in BMDM and **(J)** further quantification. Results shown are representative from 2 independent experiments. Representative microscopic images are shown from 20x magnification. Values are mean ± SEM. \*\**P* <0.01, \*\*\**P* <0.001 (LPD vs LPD+Leu).

To measure the mTOR activation in primary F4/80^+^ macrophages[50] cells were FACS-sorted from the bone marrow derived from LPD or LPD+Leu fed animals. Isolated cells were rested at 37°C for 90 minutes and stimulated with LPS (100ng/ml) for 60 minutes prior fixation, permeabilization, staining and FACS analysis of mTOR activation. We observed an increased pS6K level in LPD+Leu compared to LPD in purified F4/80^+^ cells, supporting the restoration of mTOR activation in macrophages in response to leucine supplementation of LPD *in vivo* (Fig. 5B).

In this context, we determined the effects of leucine restoration in the liver response to infection. Thus, LPD and LPD+Leu mice were infected with *S.* Typhimurium and tissues were collected 3 days later. Analysis of the liver enzymes AST and ALT in serum showed an increase in the LPD+Leu group compared to LPD, pointing to worsening in liver damage (Fig. 5C). Histological analysis of liver sections confirmed the increased liver injury in LPD+Leu mice that presented larger areas of necrosis compared to LPD fed mice after *S.* Typhimurium infection (Fig. 5D).

To further investigate the mechanisms mediating the loss of the protective effects of a LPD against infection after leucine supplementation *in vitro*, we performed in vitro experiments in BMDM cultured with in a low-aa media supplemented with leucine (Low-aa+Leu) and stimulated with LPS (Fig. 5E). Following our observations *in vivo*, in macrophages isolated from LPD+Leu mice, the analysis of mTOR pathway activation confirmed increased S6 kinase phosphorylation in BMDM cells cultured in low-aa+Leu media compared to control and low-aa media at basal conditions. S6 kinase phosphorylation was further increased upon LPS stimulation, while it remained unaffected in BMDM on Low-aa media (Fig. 5F).

To evaluate the level of autophagy, BMDM were stimulated with LPS for 2 hours, fixed, permeabilized and stained with LC3. Quantification of the number of LC3 puncta was highest in low-aa media-BMDM and the lowest in control cells (Fig. 5G). Macrophages programmed with Low-aa+Leu showed increased number of LC3 puncta compared to control, however this was still lower than Low-aa media, suggesting partial reduction of the Low-aa induced autophagy activation (Fig. 5H).

Autophagy and phagocytosis are important mechanisms regulating innate immunity. Phagocytic potential of macrophages was measured using E. coli pH Rodo bioparticles[51]. We did not observe differences in non-stimulated cells cultured with the different media, however after LPS stimulation we found a significant increase in the number of bioparticles per macrophage in Low-aa programmed cells compared to control or Low-aa+Leu cells (Fig. 5I). The Low-aa+Leu phagocytic cell potential resembled the pattern observed in control macrophages, where up to 70% of cells have 0-5 bioparticles/cell and Low-aa+Leu had 60% while in Low-aa programmed BMDM this constituted only 30% of the cells. Only 8% or 18% of cells had above 16 bioparticles per macrophage in control or Low-aa+Leu media respectively compared to 30% in Low-aa (Fig. 5J).

Finally, colocalization of red bioparticles with stained LC3 was especially striking in Low-aa programmed macrophages suggesting that Low-aa programming likely increases the phagocytic potential of BMDM, supporting that this process depends on the balance between mTOR activation, autophagy and phagocytosis[21].

Overall, amino acid stress imposed by a Low-aa shifts the balance from mTOR activation, inducing expression of proinflammatory cytokines, towards increased autophagy and bacteria phagocytosis that was reverted after mTOR restoration with Leucine supplementation. Importantly, here we demonstrate how changes in dietary protein and specific amino acid content modulate the innate immune response to infection.

## DISCUSSION

Predominantly in western countries, dietary habits are increasingly shifting towards healthier choices including the reduction of meat (hence dietary protein) intake. Still, the impact of these dietary changes, particularly in the immune response, remain largely undefined.

Here, we have studied the effects of a reduction in the protein intake (6% protein diet; LPD) rather than malnutrition, where the drastically reduced protein content (0.5-2.5% protein) causes severe impairment of the immune system function and increased susceptibility to infections[52]. In our study, we have identified the protective effect of a low protein feeding (LPD) in adult mice and the underlying molecular mechanism involving the inhibition of mTOR pathway, increased autophagy and phagocytosis in macrophages.

Gene expression analysis in total liver lysates using bulk RNA sequencing revealed the significantly decreased expression of proinflammatory cytokines, inflammasome subunits and pattern recognition receptors in LPD-fed mice compared to chow-fed mice after *S.* Typhimurium infection. Interestingly, we observed modulation of key metabolic pathways, including gluconeogenesis and lipid metabolism as well as the upregulation of Sirt3 in LPD-fed animals, a key activator of autophagy[34] and a negative regulator of the formation of the inflammasome in macrophages[35]. These results supported both the immunoregulatory and metabolic rewiring effects of the LPD intervention in response to infection that associated with a protection from Salmonella-induced liver injury.

Macrophages are a key component of the host innate immune defence against *S.* Typhimurium infection via pathogen phagocytosis and production of proinflammatory cytokines (i.e. IL1β)[53]. Previous studies showed that macrophages under protein-energy malnutrition had decreased bacteria phagocytosis and killing capacity, reduced adhesion, spreading, and fungicidal activity[54, 55], supporting that severe protein-energy malnutrition significantly impairs macrophage function and diminishes response to acute and chronic infections[55].

To pinpoint the modulation of immune cell function and more specifically of macrophages in response to LPD feeding during infection, we performed 10x Single cell RNA sequencing in immune cells isolated from mouse livers. Detailed analyses of cell-specific genetic changes using this approach evidenced decreased expression of key markers of macrophage activation like S100a8 that promotes the recruitment of leukocytes and thus a proinflammatory cytokine milieu[40]. In murine models the blockade of the S100a8/S100a9 complex with small molecules or antibodies improves pathological conditions, while decreased expression of this complex correlates with better prognosis, as sepsis surviving patients shown decreased S100A8/A9 levels compared with non-survivors[56]. Targeting S100A8/A9 can also prevent liver injury as well as bacterial dissemination at an early phase during human sepsis and endotoxemia[57].

We observed that the protection against Salmonella infection after LPD also correlated with a significantly increase in c-fos expression in monocytes, which is consistent with recent studies where deletion of *c-Fos* in mouse monocyte and macrophages led to significantly enhanced production of TNFα, IL6 and IL12 p40 in response to LPS and an increased susceptibility to *S.* Typhimurium infection[42, 43].

Interestingly, gene expression changes in monocytes were not restricted to inflammation but we also found the regulation of key metabolic factors in infected mice fed with LPD compared to control diet, including a decrease of Nupr1 and EIF1, both mediators of mTOR activation[44, 46], in monocyte/macrophages suggesting decreased activity of mTOR pathway while enhanced autophagy in these cells after LPD feeding.

The coordinated expression pattern of proinflammatory and mTOR-related genes strongly suggests mTOR role as a molecular switch and transcription factor executing nutritional and molecular programs. ChIP-sequencing datasets revealed that mTOR directly binds to thousands of regulatory regions of polymerase II-transcribed genes in both mouse liver and human prostate cancer cells[58, 59]. Interestingly, treatment of prostate cancer cells with the inhibitor of polymerase II transcriptional activity α-amanitin, which has no effect on classic cytoplasmic mTOR signaling-autophagy and phagocytosis, abrogated the metabolic reprogramming associated with the transcriptional function of nuclear mTOR observed in these cells[58]. Thus, future studies will be necessary to unravel mTOR function as transcription factor in response to a range of diets fed and at different developmental stages.

mTOR is a known regulator of macrophage activation by inhibiting autophagy[60], while autophagy is essential to control the host response to pathogens[61] via modulation of the inflammasome and IL1β production[62], e.g. impaired autophagy enhances the inflammasome activity and IL1β production in macrophages after LPS [63]. Thus, the immunomodulation and metabolic rewiring in monocytes upon LPD we observe could lead to improved ability to resolve *S.* Typhimurium infection and thus restricting liver tissue damage via modulating the mTOR/autophagy axis.

To test the regulation of the mTOR/autophagy as a mechanistic mediator of the protective effects against infection we observed upon LPD feeding *in vivo*, we performed studies *in vitro* using BMDM. Our results show that low amino acid content in culture media, mimicking low protein feeding *in vivo*, reduced macrophage proinflammatory nature and concurrently enhancing autophagy. Our results showing that reduced availability of aa in culture led to significant reduction of inflammasome-IL1β-Hif1α are in line with previous reports showing Hif1α-induced activation of the inflammasome, IL1β production and impaired autophagy flux in macrophages in patients with chronic liver inflammation during NASH[64].

Ultimately, to pinpoint the attenuation of mTOR signalling as the mechanistic mediator of the protective effects of LPD during infection we restored mTOR signaling in LPD fed mice *in vivo* by dietary supplementation of Leucine; an essential amino acid that directly activates the mTORC1 subunit of the mTOR complex[49]. In agreement with our hypothesis, leucine supplementation reverted the protective effects of a LPD in response to *S.* Typhimurium infection *in vivo*. The direct activation of mTOR by leucine-supplementation in cultured BMDM diminished the autophagy and phagocytic potential of these cells supporting the crucial immunomodulatory role of dietary amino acids, and more specifically of leucine. Our results support the reported detrimental effects of dietary-mediated mTOR activation by leucine-rich (western) diets associated with type 2 diabetes and obesity where increased leucine consumption contributes to aggravation of disease progression[65]. Obesity associates with dysfunction of innate immune response[66] and is a known risk factor for bacteraemia[67] and thus dietary interventions involving the reduction of leucine may pose beneficial potential to boost innate immunity.

Overall, our findings highlight the prospects to improve the immune response to infection using dietary interventions. Still, to fully utilise these benefits, future work is guaranteed to better understand the kinetics of metabolic changes induced by LPD to choose the most optimal age/time window.

## MATERIALS AND METHODS

### Experimental procedures in animals

C57BL/6 J mice (CD45.2), were purchased (Charles River Massachusetts, United States) and housed at the Disease Modelling Unit (University of East Anglia, Norwich, United Kingdom). All experiments were approved by the Animal Welfare and Ethical Review Body (University of East Anglia). All procedures were carried out following the guidelines of the National Academy of Sciences (National Institutes of Health, publication 86-23, revised 1985) and were performed within the provisions of the Animals (Scientific Procedures) Act 1986. Mice were kept in individually ventilated cages and housed under specific pathogen-free conditions in a 12/12-hour light/dark cycle. Animals were put on *ad libitum*, isocaloric control (Special Diets Services, 801066) or 6% low protein diet (Teklad, TD220065) for 10 weeks, and body weight was regularly monitored. Low protein diet with 3% leucine was purchased from Teklad (TD.90016). Mice used were 8-10 weeks old, male animals were used. For the differentiation of bone marrow derived macrophages mice were exposed to CO_2_, BM was harvested for in vitro differentiation.

### *S.* Typhimurium infection model

Glycerol stock of Salmonella enterica serotype Typhimurium (SL1344-JH3009) was plated on Luria Broth agar plates and the colonies were inoculated and grown overnight into 5 ml of Luria Broth with 0.3 M NaCl (LBS). The overnight culture was then diluted 1:100 in LBS and grown until the culture optical density (ΔOD600nm) of 1.2 – 1.4 (late exponential phase). This is the time point where SPI1 invasion genes are turned on in *S.* Typhimurium. The bacterial culture was then centrifuged at 3000 × g for 7 minutes before washing bacterial cells twice in 25 ml of sterile DPBS at room temperature. Finally, resuspend the bacterial cells in sterile DPBS at a concentration of 1–5 × 10^8^ CFU per 100 μl of DPBS (knowing that DOD600nm 1.26 corresponds to 7.53 × 10^8^ CFU/ml). Mice were infected with 100 μl of 1 × 10^8^ CFU *S.* Typhimurium (SL1344-JH3009) by intraperitoneal injection for 3 days. The mice were anesthetized using isoflurane to collect blood, followed by exposure to CO2, and the liver was collected for flow cytometric analysis, sorting and sequencing.

### Liver histology

Liver tissues were harvested and immediately fixed in 10% neutral formalin and embedded in paraffin blocks 24 hours later. Tissue blocks were sectioned, dewaxed, and hydrated prior to being stained with Hematoxylin & Eosin (H&E) for histopathological analysis. Slides were imaged using brightfield on a BX53 upright microscope (Olympus) with an Olympus DP74 colour camera and a pT100 LED transmitted light source (CoolLED).

### Serum transaminases

The levels of circulating ALT and AST were measured in serum samples in a Randox RX Daytona analyser.

### BMDM Differentiation and culture

BMDMs were differentiated from bone marrow cells isolated from WT mice. The femur and tibia were cut in the middle and placed in a 0.5 ml Eppendorf tube in which a hole was made to allow the removal of the BM, placed in an intact 1.5 ml Eppendorf and centrifuged 1000 × g for 6 seconds to collect the BM cells. The BM pellet from each mouse was pooled and plated with RPMI-1640 (Gibco, Thermo Fisher Scientific) supplemented with 20% foetal bovine serum (FBS) (Gibco, Thermo Fisher Scientific), 30% L929 conditioned media and 1% penicillin/streptomycin (Gibco, Thermo Fisher Scientific). Cells were allowed to differentiate for 7 days, with fresh media added on day 3. A total of 1 x 10e6 adherent cells were then plated for experiments in 6-well plates and 500k for 12-well plates. For immunofluorescence imaging of BMDMs, cells were plated on glass coverslips.

BMDMs were then cultured in DMEM-LM medium (Thermo scientific), supplemented with 3% L929 conditioned media and 1% penicillin/streptomycin. To replicate normal diet, 10% FBS and 1 X MEM amino acid solution (Gibco, ThermoFisher Scientific) were also added to the medium, and to replicate LPD diet in mice, a low amino acid media (low-aa) was created by adding 1% FBS and 0.2 X MEM amino acid solution (adjusted for the leucine concentration) for 48 hours. To assess the effect of leucine in low-aa medium, 1965mg/L leucine and 1 X glucose (Gibco, ThermoFisher Scientific) was added to low-aa medium. BMDMs were then starved by removing FBS from the mediums for 12 hours prior to 100ng/mL LPS stimulation.

### Assessment of LC3-associated autophagy

For determination of autophagy, all cells were pretreated with 20 mM NH4Cl and 100 mM leupeptin to inhibit lysosomal proteolysis, 2 hours before LPS treatment. Cells were then fixed using 4% formaldehyde solution, buffered pH 6.9 (Sigma-Aldrich) and permeabilised using solution B from FIX & PERM^TM^ Cell Permeabilization Kit (Invitrogen^TM^, ThermoFisher scientific), during which cells were stained with 1:300 dilution of LC3-FITC antibody, EPR18709 (Abcam). Cells were washed and mounted using Vectashield Antifade mounting medium with DAPI (Vector Labs). imaged using AxioImager M2 (Zeiss) using x63 magnification oil immersion. LC3-positive areas per macrophage were quantified using Fiji Image J (2.9.0/1.53t).

### Quantitative Real-Time PCR

RNA from cells was isolated using the ReliaPrep RNA Miniprep System (Promega). RNA was reverse transcribed using UltraScript^TM^ cDNA Synthesis Kit (PCR Biosystems) and qRT-PCR was performed using qPCRBIO SyGreen Mix (PCR Biosystems). Primer sequences can be made available upon request.

### Flow Cytometry and sorting

Immune cells were isolated from the mouse liver, as described previously[68, 69]. Immune cells were stained with CD45-APC-Cy7 (Becton Dickinson, Franklin Lakes, NJ), CD11b-PE (Becton Dickinson), F4/80-FITC (Miltenyi Biotec), and Ly6C-Pacific blue (MACS). Flow cytometry was carried out using BD LSR-Fortessa. Analysis was performed using FlowJo software (FlowJo 10.8.1, Ashland, OR). For single cell RNA seq cells were stained with CD45-APC-Cy7, and DAPI, and sorted on or BD Aria-Fusion to purify viable, CD45+ cells that were loaded on 10X.

For intracellular FACS staining metabolically reprogrammed BMDMs were dissociated from cell culture plates and fixed using ice cold methanol, washed and stained with 0.1 uL ps6k, cupk43k (eBioscience^TM^) per sample. A BD FACS Symphony A1 (Becton, Dickinson and Company), was used to assess pS6K expression and data was analysed using FlowJo 10.9.0 (Becton, Dickinson and Company).

### Sequencing of single-cell cDNA libraries

Sorted cells were processed by 3’ end single-cell RNA-Seq using the 10X Genomics Chromium (V3 Kit) according to the manufacturer’s protocol (10X Genomics, Pleasanton, CA). Libraries were sequenced on a NovaSeq 6000 (Illumina, San Diego) in paired-end, single index mode as per the 10X Genomics recommended metrics.

The single cell data was processed using 10X Genomics Cell Ranger analysis pipeline (cellranger-6.0.1) with Ensembl GRCm39 Mus musculus assembly and gene annotation. A feature barcode matrix was generated for each sample by applying the *cellranger count* pipeline. All feature barcode matrices were aggregated using *cellranger aggr*, which normalises sequencing depth across samples. QC was performed excluding cells with fewer than 1000 genes detected or more than 5% of UMI counts associated to mitochondrial genes. In total, 7,148 cells were selected, distributed as follows: 3,523 cells came from the immune liver cells S.Typhimurium treated Control diet sample and 3, 625 cells from the immune liver cells S.Typhimurium treated Low Protein diet sample. Cell cycle variation was removed using the ‘CellCycleScoring’ method followed by regressing out ‘S.Score’ and ‘G2M.Score’. Cells were then normalized to 10,000 UMIs per cell and logarithmically transformed. HVGs were selected using the ‘FindVariableFeatures’ method. UMAP visualizations were obtained from 20 PCA components, and clusters were defined at a resolution of 0.3 using Louvain algorithm. Cell types were annotated using typical marker genes for the different haematopoietic populations. Differential gene expression was performed using the ‘FindMarkers’ method.

### Statistical analysis

Statistical analyses were performed using GraphPad Prism software version 10.0.3. Statistical differences between two groups were determined by unpaired, two-tailed Student’s t-test with Welch’s correction.

For Bulk-RNAseq on liver tissues, q-values were determined through Benjamini-Hochberg p-value adjustment and q-value were <0.001. For single cell RNA-seq adjusted p-values were derived using Benjamini-Hochberg multiple test correction.

Two-way ANOVA tests were used when comparing different treatments at different timepoints and were performed using GraphPad Prism software settings. Data are shown as mean±SEM. *p<0.033, **p<0.002, ***p<0.001.

## Data availability statement

The authors declare that all data generated from this study are available within the manuscript and the supplemental material provided. Any additional files or information can be provided upon request to the corresponding authors.

## Acknowledgments

The authors gratefully acknowledge the support of the Biotechnology and Biological Sciences Research Council (BBSRC) Institute Strategic Programme Gut Microbes and Health BB/R012490/1 and its constituent project BBS/E/F/000PR10355, and the BBSRC Core Capability Grant BB/CCG1860/1 as well as the BBSRC Institute Strategic Programme Food Microbiomes and Health BB/X011054/1 and its constituent project BBS/E/F/000PR13632 (NB) and Microbes and Food Safety (BB/X011011/1) And its constituent BBS/E/F/000PR13634. KH was supported by the UKRI BBSRC Norwich research park Bioscience doctoral training programme BB/T008717/1. PR is supported by a BBSRC response mode BB/W002450/1 (to NB). SAR was supported by the UKRI MRC project (MR/T02934X/1). CH was supported by the Wellcome Trust Clinical Research Fellowship (220534/Z/20/Z). EW and ICM acknowledge support from the Biotechnology and Biological Sciences Research Council (BBSRC), part of UK Research and Innovation, Core Capability Grant BB/CCG1720/1 and the National Capability BBS/E/T/000PR9816, BBS/E/T/000PR9814, BBS/E/T/000PR9811. EW and ICM acknowledge the support of the Biotechnology and Biological Sciences Research Council (BBSRC), part of UK Research and Innovation; Earlham Institute Strategic Programme Grant Cellular Genomics BBX011070/1 and its constituent work packages(s) - choose one or more of the three CellGen work packages we contribute to below: BBS/E/ER/230001B and BBS/E/ER/230001C. EW was supported by the Sir Henry Wellcome Postdoctoral fellowship (213731/Z/18/Z). ICM was additionally supported by BBSRC New Investigator Grant BB/P022073/1.

We thank Dr W. Haerty for insightful discussions and critical comments on the manuscript.

## REFERENCES

[1] Childs CE, Calder PC, Miles EA. Diet and Immune Function. Nutrients 2019;11(8).

[2] Schulze MB, Martinez-Gonzalez MA, Fung TT, Lichtenstein AH, Forouhi NG. Food based dietary patterns and chronic disease prevention. BMJ 2018;361:k2396.

[3] Christ A, Gunther P, Lauterbach MAR, Duewell P, Biswas D, Pelka K, Scholz CJ, Oosting M, Haendler K, Bassler K, Klee K, Schulte-Schrepping J, Ulas T, Moorlag S, Kumar V, Park MH, Joosten LAB, Groh LA, Riksen NP, Espevik T, Schlitzer A, Li Y, Fitzgerald ML, Netea MG, Schultze JL, Latz E. Western Diet Triggers NLRP3-Dependent Innate Immune Reprogramming. Cell 2018;172(1-2):162–75 e14.

[4] Cildir G, Akincilar SC, Tergaonkar V. Chronic adipose tissue inflammation: all immune cells on the stage. Trends Mol Med 2013;19(8):487–500.

[5] Netea MG, Joosten LA, Latz E, Mills KH, Natoli G, Stunnenberg HG, O’Neill LA, Xavier RJ. Trained immunity: A program of innate immune memory in health and disease. Science 2016;352(6284):aaf1098.

[6] Kaspersen KA, Pedersen OB, Petersen MS, Hjalgrim H, Rostgaard K, Moller BK, Juul-Sorensen C, Kotze S, Dinh KM, Erikstrup LT, Sorensen E, Thorner LW, Burgdorf KS, Ullum H, Erikstrup C. Obesity and risk of infection: results from the Danish Blood Donor Study. Epidemiology 2015;26(4):580–9.

[7] Maccioni L, Weber S, Elgizouli M, Stoehlker AS, Geist I, Peter HH, Vach W, Nieters A. Obesity and risk of respiratory tract infections: results of an infection-diary based cohort study. BMC Public Health 2018;18(1):271.

[8] Mariotti F, Gardner CD. Dietary Protein and Amino Acids in Vegetarian Diets-A Review. Nutrients 2019;11(11).

[9] Brown EM, Wlodarska M, Willing BP, Vonaesch P, Han J, Reynolds LA, Arrieta MC, Uhrig M, Scholz R, Partida O, Borchers CH, Sansonetti PJ, Finlay BB. Diet and specific microbial exposure trigger features of environmental enteropathy in a novel murine model. Nat Commun 2015;6:7806.

[10] Onyango AW, Jean-Baptiste J, Samburu B, Mahlangu TLM. Regional Overview on the Double Burden of Malnutrition and Examples of Program and Policy Responses: African Region. Ann Nutr Metab 2019;75(2):127–30.

[11] Rivadeneira DE, Grobmyer SR, Naama HA, Mackrell PJ, Mestre JR, Stapleton PP, Daly JM. Malnutrition-induced macrophage apoptosis. Surgery 2001;129(5):617–25.

[12] Salameh E, Morel FB, Zeilani M, Dechelotte P, Marion-Letellier R. Animal Models of Undernutrition and Enteropathy as Tools for Assessment of Nutritional Intervention. Nutrients 2019;11(9).

[13] Fontana L, Adelaiye RM, Rastelli AL, Miles KM, Ciamporcero E, Longo VD, Nguyen H, Vessella R, Pili R. Dietary protein restriction inhibits tumor growth in human xenograft models. Oncotarget 2013;4(12):2451–61.

[14] Levine ME, Suarez JA, Brandhorst S, Balasubramanian P, Cheng CW, Madia F, Fontana L, Mirisola MG, Guevara-Aguirre J, Wan J, Passarino G, Kennedy BK, Wei M, Cohen P, Crimmins EM, Longo VD. Low protein intake is associated with a major reduction in IGF-1, cancer, and overall mortality in the 65 and younger but not older population. Cell Metab 2014;19(3):407–17.

[15] Solanki S, Sanchez K, Ponnusamy V, Kota V, Bell HN, Cho CS, Kowalsky AH, Green M, Lee JH, Shah YM. Dysregulated Amino Acid Sensing Drives Colorectal Cancer Growth and Metabolic Reprogramming Leading to Chemoresistance. Gastroenterology 2023;164(3):376–91 e13.

[16] Ginhoux F, Guilliams M. Tissue-Resident Macrophage Ontogeny and Homeostasis. Immunity 2016;44(3):439–49.

[17] Arango Duque G, Descoteaux A. Macrophage cytokines: involvement in immunity and infectious diseases. Front Immunol 2014;5:491.

[18] Hirayama D, Iida T, Nakase H. The Phagocytic Function of Macrophage-Enforcing Innate Immunity and Tissue Homeostasis. Int J Mol Sci 2017;19(1).

[19] Ilyas B, Tsai CN, Coombes BK. Evolution of Salmonella-Host Cell Interactions through a Dynamic Bacterial Genome. Front Cell Infect Microbiol 2017;7:428.

[20] Robinson N, McComb S, Mulligan R, Dudani R, Krishnan L, Sad S. Type I interferon induces necroptosis in macrophages during infection with Salmonella enterica serovar Typhimurium. Nat Immunol 2012;13(10):954–62.

[21] Ganesan R, Hos NJ, Gutierrez S, Fischer J, Stepek JM, Daglidu E, Kronke M, Robinson N. Salmonella Typhimurium disrupts Sirt1/AMPK checkpoint control of mTOR to impair autophagy. PLoS Pathog 2017;13(2):e1006227.

[22] Giovannini L, Bianchi S. Role of nutraceutical SIRT1 modulators in AMPK and mTOR pathway: Evidence of a synergistic effect. Nutrition 2017;34:82–96.

[23] Weichhart T, Hengstschlager M, Linke M. Regulation of innate immune cell function by mTOR. Nat Rev Immunol 2015;15(10):599–614.

[24] Orecchioni M, Ghosheh Y, Pramod AB, Ley K. Macrophage Polarization: Different Gene Signatures in M1(LPS+) vs. Classically and M2(LPS-) vs. Alternatively Activated Macrophages. Front Immunol 2019;10:1084.

[25] Palmieri EM, Gonzalez-Cotto M, Baseler WA, Davies LC, Ghesquiere B, Maio N, Rice CM, Rouault TA, Cassel T, Higashi RM, Lane AN, Fan TW, Wink DA, McVicar DW. Nitric oxide orchestrates metabolic rewiring in M1 macrophages by targeting aconitase 2 and pyruvate dehydrogenase. Nat Commun 2020;11(1):698.

[26] Palsson-McDermott EM, Curtis AM, Goel G, Lauterbach MAR, Sheedy FJ, Gleeson LE, van den Bosch MWM, Quinn SR, Domingo-Fernandez R, Johnston DGW, Jiang JK, Israelsen WJ, Keane J, Thomas C, Clish C, Vander Heiden M, Xavier RJ, O’Neill LAJ. Pyruvate Kinase M2 Regulates Hif-1alpha Activity and IL-1beta Induction and Is a Critical Determinant of the Warburg Effect in LPS-Activated Macrophages. Cell Metab 2015;21(2):347.

[27] Wallace C, Keast D. Glutamine and macrophage function. Metabolism 1992;41(9):1016–20.

[28] Rodriguez AE, Ducker GS, Billingham LK, Martinez CA, Mainolfi N, Suri V, Friedman A, Manfredi MG, Weinberg SE, Rabinowitz JD, Chandel NS. Serine Metabolism Supports Macrophage IL-1beta Production. Cell Metab 2019;29(4):1003–11 e4.

[29] Ellyard JI, Quah BJ, Simson L, Parish CR. Alternatively activated macrophage possess antitumor cytotoxicity that is induced by IL-4 and mediated by arginase-1. J Immunother 2010;33(5):443–52.

[30] Son SM, Park SJ, Lee H, Siddiqi F, Lee JE, Menzies FM, Rubinsztein DC. Leucine Signals to mTORC1 via Its Metabolite Acetyl-Coenzyme A. Cell Metab 2019;29(1):192–201 e7.

[31] Cunningham JT, Moreno MV, Lodi A, Ronen SM, Ruggero D. Protein and nucleotide biosynthesis are coupled by a single rate-limiting enzyme, PRPS2, to drive cancer. Cell 2014;157(5):1088–103.

[32] Huang J, Brumell JH. Bacteria-autophagy interplay: a battle for survival. Nat Rev Microbiol 2014;12(2):101–14.

[33] Yoon BR, Oh YJ, Kang SW, Lee EB, Lee WW. Role of SLC7A5 in Metabolic Reprogramming of Human Monocyte/Macrophage Immune Responses. Front Immunol 2018;9:53.

[34] Zhang T, Liu J, Tong Q, Lin L. SIRT3 Acts as a Positive Autophagy Regulator to Promote Lipid Mobilization in Adipocytes via Activating AMPK. Int J Mol Sci 2020;21(2).

[35] Liu P, Huang G, Wei T, Gao J, Huang C, Sun M, Zhu L, Shen W. Sirtuin 3-induced macrophage autophagy in regulating NLRP3 inflammasome activation. Biochim Biophys Acta Mol Basis Dis 2018;1864(3):764–77.

[36] Isaacs-Ten A, Moreno-Gonzalez M, Bone C, Martens A, Bernuzzi F, Ludwig T, Hellmich C, Hiller K, Rushworth SA, Beraza N. Metabolic Regulation of Macrophages by SIRT1 Determines Activation During Cholestatic Liver Disease in Mice. Cell Mol Gastroenterol Hepatol 2022;13(4):1019–39.

[37] Jia L, Shi Y, Wen Y, Li W, Feng J, Chen C. The roles of TNFAIP2 in cancers and infectious diseases. J Cell Mol Med 2018;22(11):5188–95.

[38] Schwanzer-Pfeiffer D, Rossmanith E, Schildberger A, Falkenhagen D. Characterization of SVEP1, KIAA, and SRPX2 in an in vitro cell culture model of endotoxemia. Cell Immunol 2010;263(1):65–70.

[39] Lee WB, Choi WY, Lee DH, Shim H, Kim-Ha J, Kim YJ. OAS1 and OAS3 negatively regulate the expression of chemokines and interferon-responsive genes in human macrophages. BMB Rep 2019;52(2):133–8.

[40] Pruenster M, Vogl T, Roth J, Sperandio M. S100A8/A9: From basic science to clinical application. Pharmacol Ther 2016;167:120–31.

[41] Quintero PA, Knolle MD, Cala LF, Zhuang Y, Owen CA. Matrix metalloproteinase-8 inactivates macrophage inflammatory protein-1 alpha to reduce acute lung inflammation and injury in mice. J Immunol 2010;184(3):1575–88.

[42] Maruyama K, Sano G, Ray N, Takada Y, Matsuo K. c-Fos-deficient mice are susceptible to Salmonella enterica serovar Typhimurium infection. Infect Immun 2007;75(3):1520–3.

[43] Ray N, Kuwahara M, Takada Y, Maruyama K, Kawaguchi T, Tsubone H, Ishikawa H, Matsuo K. c-Fos suppresses systemic inflammatory response to endotoxin. Int Immunol 2006;18(5):671–7.

[44] Li A, Li X, Chen X, Zeng C, Wang Z, Li Z, Chen J. NUPR1 Silencing Induces Autophagy-Mediated Apoptosis in Multiple Myeloma Cells Through the PI3K/AKT/mTOR Pathway. DNA Cell Biol 2020;39(3):368–78.

[45] Tattoli I, Sorbara MT, Vuckovic D, Ling A, Soares F, Carneiro LA, Yang C, Emili A, Philpott DJ, Girardin SE. Amino acid starvation induced by invasive bacterial pathogens triggers an innate host defense program. Cell Host Microbe 2012;11(6):563–75.

[46] Hui KK, Chen YK, Endo R, Tanaka M. Translation from the Ribosome to the Clinic: Implication in Neurological Disorders and New Perspectives from Recent Advances. Biomolecules 2019;9(11).

[47] Biasizzo M, Kopitar-Jerala N. Interplay Between NLRP3 Inflammasome and Autophagy. Front Immunol 2020;11:591803.

[48] Takahara T, Amemiya Y, Sugiyama R, Maki M, Shibata H. Amino acid-dependent control of mTORC1 signaling: a variety of regulatory modes. J Biomed Sci 2020;27(1):87.

[49] Lynch CJ. Role of leucine in the regulation of mTOR by amino acids: revelations from structure-activity studies. J Nutr 2001;131(3):861S–5S.

[50] Austyn JM, Gordon S. F4/80, a monoclonal antibody directed specifically against the mouse macrophage. Eur J Immunol 1981;11(10):805–15.

[51] Moore JA, Mistry JJ, Hellmich C, Horton RH, Wojtowicz EE, Jibril A, Jefferson M, Wileman T, Beraza N, Bowles KM, Rushworth SA. LC3-associated phagocytosis in bone marrow macrophages suppresses acute myeloid leukemia progression through STING activation. J Clin Invest 2022;132(5).

[52] Bourke CD, Berkley JA, Prendergast AJ. Immune Dysfunction as a Cause and Consequence of Malnutrition. Trends Immunol 2016;37(6):386–98.

[53] Shukla S, Telraja J, Yadav M, Prakash H. Editorial: Modulation of Macrophage Signaling Pathways During Bacterial Infections. Front Cell Infect Microbiol 2021;11:689759.

[54] Corware K, Yardley V, Mack C, Schuster S, Al-Hassi H, Herath S, Bergin P, Modolell M, Munder M, Muller I, Kropf P. Protein energy malnutrition increases arginase activity in monocytes and macrophages. Nutr Metab (Lond) 2014;11(1):51.

[55] Redmond HP, Leon P, Lieberman MD, Hofmann K, Shou J, Reynolds JV, Goldfine J, Johnston RB, Jr., Daly JM. Impaired macrophage function in severe protein-energy malnutrition. Arch Surg 1991;126(2):192–6.

[56] Payen D, Lukaszewicz AC, Belikova I, Faivre V, Gelin C, Russwurm S, Launay JM, Sevenet N. Gene profiling in human blood leucocytes during recovery from septic shock. Intensive Care Med 2008;34(8):1371–6.

[57] van Zoelen MA, Vogl T, Foell D, Van Veen SQ, van Till JW, Florquin S, Tanck MW, Wittebole X, Laterre PF, Boermeester MA, Roth J, van der Poll T. Expression and role of myeloid-related protein-14 in clinical and experimental sepsis. Am J Respir Crit Care Med 2009;180(11):1098–106.

[58] Audet-Walsh E, Dufour CR, Yee T, Zouanat FZ, Yan M, Kalloghlian G, Vernier M, Caron M, Bourque G, Scarlata E, Hamel L, Brimo F, Aprikian AG, Lapointe J, Chevalier S, Giguere V. Nuclear mTOR acts as a transcriptional integrator of the androgen signaling pathway in prostate cancer. Genes Dev 2017;31(12):1228–42.

[59] Chaveroux C, Eichner LJ, Dufour CR, Shatnawi A, Khoutorsky A, Bourque G, Sonenberg N, Giguere V. Molecular and genetic crosstalks between mTOR and ERRalpha are key determinants of rapamycin-induced nonalcoholic fatty liver. Cell Metab 2013;17(4):586–98.

[60] Ko JH, Yoon SO, Lee HJ, Oh JY. Rapamycin regulates macrophage activation by inhibiting NLRP3 inflammasome-p38 MAPK-NFkappaB pathways in autophagy- and p62-dependent manners. Oncotarget 2017;8(25):40817–31.

[61] Ma Y, Galluzzi L, Zitvogel L, Kroemer G. Autophagy and cellular immune responses. Immunity 2013;39(2):211–27.

[62] Harris J, Hartman M, Roche C, Zeng SG, O’Shea A, Sharp FA, Lambe EM, Creagh EM, Golenbock DT, Tschopp J, Kornfeld H, Fitzgerald KA, Lavelle EC. Autophagy controls IL-1beta secretion by targeting pro-IL-1beta for degradation. J Biol Chem 2011;286(11):9587–97.

[63] Saitoh T, Fujita N, Jang MH, Uematsu S, Yang BG, Satoh T, Omori H, Noda T, Yamamoto N, Komatsu M, Tanaka K, Kawai T, Tsujimura T, Takeuchi O, Yoshimori T, Akira S. Loss of the autophagy protein Atg16L1 enhances endotoxin-induced IL-1beta production. Nature 2008;456(7219):264-8.

[64] Wang X, de Carvalho Ribeiro M, Iracheta-Vellve A, Lowe P, Ambade A, Satishchandran A, Bukong T, Catalano D, Kodys K, Szabo G. Macrophage-Specific Hypoxia-Inducible Factor-1alpha Contributes to Impaired Autophagic Flux in Nonalcoholic Steatohepatitis. Hepatology 2019;69(2):545–63.

[65] Melnik BC. Leucine signaling in the pathogenesis of type 2 diabetes and obesity. World J Diabetes 2012;3(3):38–53.

[66] Huttunen R, Laine J, Lumio J, Vuento R, Syrjanen J. Obesity and smoking are factors associated with poor prognosis in patients with bacteraemia. BMC Infect Dis 2007;7:13.

[67] Pugliese G, Liccardi A, Graziadio C, Barrea L, Muscogiuri G, Colao A. Obesity and infectious diseases: pathophysiology and epidemiology of a double pandemic condition. Int J Obes (Lond) 2022;46(3):449–65.

[68] Blokker BA, Maijo M, Echeandia M, Galduroz M, Patterson AM, Ten A, Philo M, Schungel R, Gutierrez-de Juan V, Halilbasic E, Fuchs C, Le Gall G, Milkiewicz M, Milkiewicz P, Banales JM, Rushbrook SM, Mato JM, Trauner M, Muller M, Martinez-Chantar ML, Varela-Rey M, Beraza N. Fine-Tuning of Sirtuin 1 Expression Is Essential to Protect the Liver From Cholestatic Liver Disease. Hepatology 2019;69(2):699–716.

[69] Isaacs-Ten A, Echeandia M, Moreno-Gonzalez M, Brion A, Goldson A, Philo M, Patterson AM, Parker A, Galduroz M, Baker D, Rushbrook SM, Hildebrand F, Beraza N. Intestinal Microbiome-Macrophage Crosstalk Contributes to Cholestatic Liver Disease by Promoting Intestinal Permeability in Mice. Hepatology 2020;72(6):2090–108.

