## Supplemental material for "Low protein diet protects liver function upon Salmonella infection by metabolic reprogramming of macrophages"

### Supplementary Figure 1

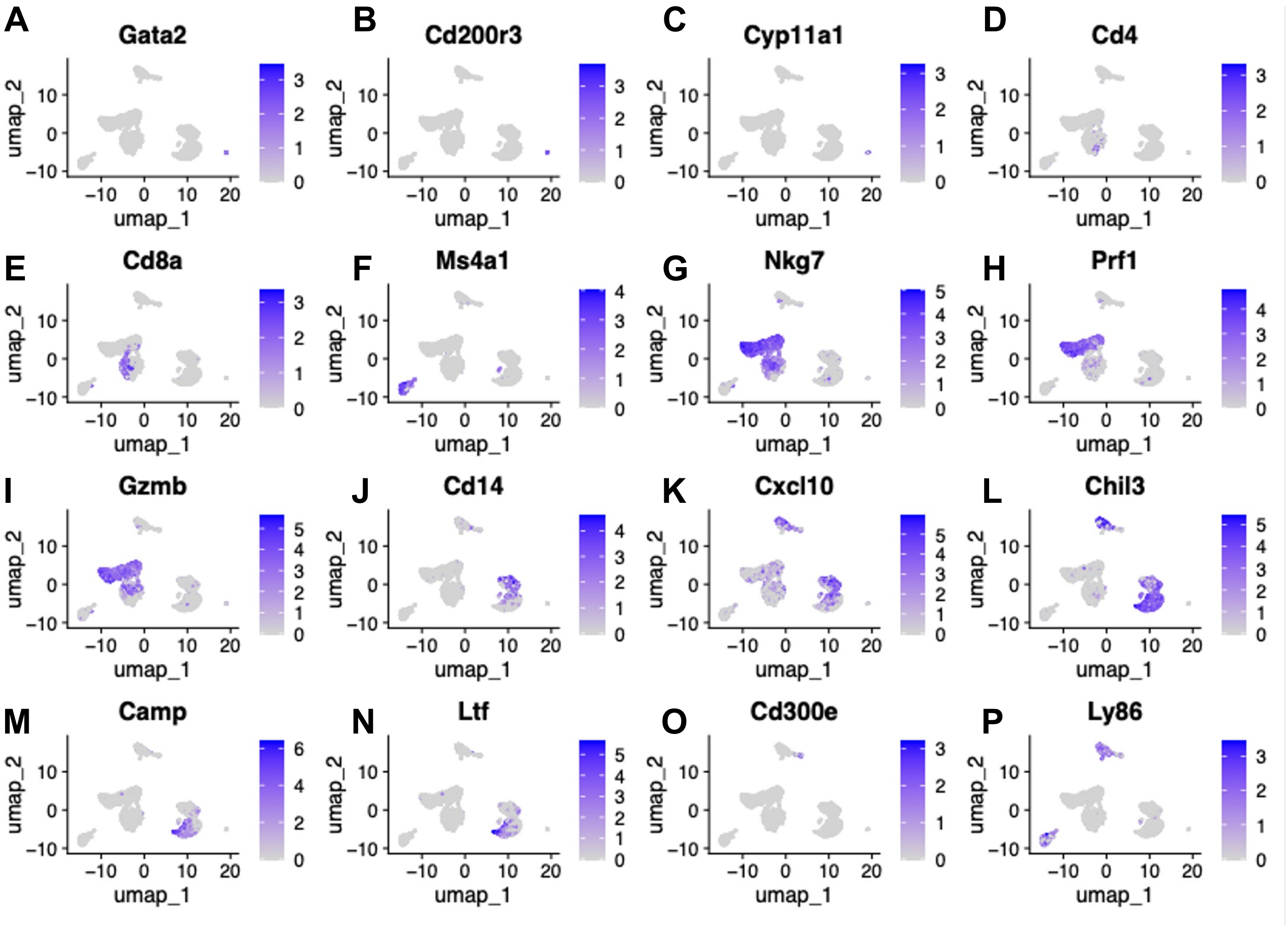


Suppl Fig 1: Feature plots depicting marker gene expression in cell clusters

### Supplementary Table 1

| Gene Symbol | Ensembl ID | Gene | Control | | LPD | |
|---|---|---|---|---|---|---|
| | | | Fold Change[a] | q-value[b] | Fold Change[a] | q-value[b] |
| **Ccl2** | ENSMUSG00000035385 | Chemokine (C-C motif) ligand 2 | 5.4 | <0.001 | 4.7 | <0.001 |
| **Ccl5** | ENSMUSG00000035042 | Chemokine (C-C motif) ligand 5 | 5.2 | <0.001 | 2.7 | <0.001 |
| **Cd14** | ENSMUSG00000051439 | Cluster of differentiation 14 | 7.7 | <0.001 | 6.1 | <0.001 |
| **Cxcl2** | ENSMUSG00000058427 | Chemokine (C-X-C motif) ligand 2 | 6.9 | <0.001 | 5.3 | <0.001 |
| **Cxcl9** | ENSMUSG00000029417 | Chemokine (C-X-C motif) ligand 9 | 9.4 | <0.001 | 7.5 | <0.001 |
| **Cxcl10** | ENSMUSG00000034855 | Chemokine (C-X-C motif) ligand 10 | 7.4 | <0.001 | 3.4 | <0.001 |
| **Ifng** | ENSMUSG00000055170 | Interferon Gamma | 8.2 | <0.001 | 7.5 | <0.001 |
| **Il1b** | ENSMUSG00000027398 | Interleukin 1 beta | 5.2 | <0.001 | 3.6 | <0.001 |
| **Il6** | ENSMUSG00000025746 | Interleukin 6 | 7.5 | <0.001 | 5.2 | <0.001 |
| **Il12a** | ENSMUSG00000027776 | Interleukin 12a | 5.3 | <0.001 | 3.0 | <0.001 |
| **Il12b** | ENSMUSG00000004296 | Interleukin 12b | 8.0 | <0.001 | 5.5 | <0.001 |
| **S100a9** | ENSMUSG00000056071 | S100 calcium-binding protein A9 | 7.0 | <0.001 | 4.8 | <0.001 |
| **Tlr2** | ENSMUSG00000027995 | Toll-like receptor 2 | 5.8 | <0.001 | 3.8 | <0.001 |
| **Tnf** | ENSMUSG00000024401 | Tumor necrosis factor | 7.1 | <0.001 | 5.1 | <0.001 |
| **Trem1** | ENSMUSG00000042265 | Triggering receptor expressed on myeloid cells 1 | 6.4 | <0.001 | 5.0 | <0.001 |

[a] Fold change of Salmonella compared to Control.

[b]q-values were determined through Benjamini-Hochberg p-value adjustment.

**Supplementary Table 1: Bulk RNA-seq gene enrichment analysis indicated gene changes in liver cells compared to control**

**Supplementary Fig 1: Feature plot depicting marker gene expression in cell clusters** **(A-C)** Basophiles basophils: *Cd200r3*, *Cyp11a1*, *Gata2,* **(D)** CD4 T cells, **(E)** Cd8+ T cells, **(F)** Ms4a1 B cells, **(G-I)** mature NK cells**, (J-K)** monocytes**, (L-N)** neutrophils, **(O-P)** Dendritic cells.
